# Using physiological signals to predict temporal defense responses: a multi-modality analysis

**DOI:** 10.1101/2020.12.17.423337

**Authors:** Tae-jun Choi, Honggu Lee

## Abstract

Defense responses are a highly conserved behavioral response set across species. Defense responses motivate organisms to detect and react to threats and potential danger as a precursor to anxiety. Accurate measurement of temporal defense responses is important for understanding clinical anxiety and mood disorders, such as post-traumatic stress disorder, obsessive compulsive disorder, and generalized anxiety disorder. Within these conditions, anxiety is defined as a state of prolonged defense response elicitation to a threat that is ambiguous or unspecific. In this study, we aimed to develop a data-driven approach to capture temporal defense response elicitation through a multi-modality data analysis of physiological signals, including electroencephalogram (EEG), electrocardiogram (ECG), and eye-tracking information. A fear conditioning paradigm was adopted to develop a defense response classification model. From a classification model based on 42 feature sets, a higher order crossing feature set-based model was chosen for further analysis with cross-validation loss of 0.0462 (SEM: 0.0077). To validate our model, we compared predicted defense response occurrence ratios from a comprehensive situation that generates defense responses by watching movie clips with fear awareness and threat existence predictability, which have been reported to correlate with defense response elicitation in previous studies. We observed that defense response occurrence ratios are correlated with threat existence predictability, but not with fear awareness. These results are similar to those of previous studies using comprehensive situations. Our study provides insight into measurement of temporal defense responses via a novel approach, which can improve understanding of anxiety and related clinical disorders for neurobiological and clinical researchers.

## Introduction

Defense responses are a set of responses to threat stimuli that have evolved to protect against threats and reduce harms for threatened organisms [1]. Defense responses are a highly conserved behavior across species that develops not only through learning from experience, but also by evolution [2, 3]. Defense responses, such as flight, avoidance, freezing, and defensive threat, are observed in variouss species and occur quickly in response to threat-related stimuli or situations. Moreover, defense responses are an important component of clinical anxiety and mood disorders, such as post-traumatic stress disorder (PTSD), obsessive compulsive disorder (OCD), and generalized anxiety disorder (GAD) [4–6]. Within these, anxiety is defined as a state of prolonged defense response elicitation to a threat that is ambiguous or unspecific [7]. Therefore, accurate measurement of defense responses is an important issue for understanding clinical anxiety and related disorders.

Recent studies of emotions associated with defense responses have focused on emotions that induce a defense response and have measured these with various indicators. Meanwhile, modulation of defense responses has mainly been performed with the fear conditioning paradigm. Fear conditioning (also known as threat conditioning [4]) is a type of Pavlovian classical conditioning and is one of the most extensively used methods for investigating fear and anxiety behaviors in both psychology and neurobiology. Beneficial features of fear conditioning include its simplicity, rapid acquisition, and excellent cross-species translational value from rodents to humans [8]. Fear conditioning produces a pairing between a neutral stimulus (conditioned stimulus, CS+) and an aversive stimulus (unconditioned stimulus, US). Once sufficient contingency between the CS+ and US has been established, behaviors or physiological responses, for example defense responses, that are elicited by the CS+ are known as conditioned responses (CRs).

In animal studies, defense responses are measured by observing behaviors such as freezing and flight [9, 10]. In contrast, human studies measure defense responses explicitly through self-report or implicitly by measuring fear-potentiated startle response (FPS) or physiological signals such as heartbeat, skin conductance, and pupil dilation [11–15]. Self-report is most commonly used because of the low cost and ease of implementation. However, the decision-making process that occurs to answer a question may prove problematic as it may lead to inaccurate results that differ from what the participant experiences in an actual experiment. Furthermore, self-report is subject to differences in individual perception and intentional distortion of questions.

For these reasons, many conditioning studies measure defense responses with FPS. FPS is a skeletal musculature contraction reflex that is elicited during defense responses produced by unexpected or threatening sensory events. It is mediated by the brainstem and amygdala [16–21] and is generally evoked by a loud noise. FPS provides an implicit measure of fear represented by the magnitude of startle reactivity at a certain point in time measured with electrooculogram. Therefore, it is difficult to measure the frequency of defense responses with FPS in an environment with unpredictable threats. Moreover, a previous study reports that using FPS affects immersiveness of a study and may itself influence fear acquisition [22]. Physiological signals, such as heartbeat, skin conductance, and pupil dilations can also be used as measures of defense response. However, these are nonspecific measures of arousal that cannot easily be interpreted as representing emotional reactions. Moreover, these signals are not suited for measuring continuous elicitation of defense responses due to slow reaction and recovery speeds in response to stimuli.

In this study, the aim was to develop a new way to capture defense responses with a multi-modality data analysis that uses physiological signals from electroencephalogram (EEG), electrocardiogram (ECG), and eye-tracking data. Previous studies have reported the characteristics of EEG during defense responses. In an animal study, 4 Hz oscillation was observed in the prefrontal cortex during freezing [10]. Furthermore, in humans, an oscillatory power change was observed at different band frequencies in the prefrontal, frontal, and midline cortices [23]. In particular, EEG has the advantage of capturing neural activity underlying defense responses due to its excellent temporal resolution. Therefore, EEG was measured in the prefrontal cortex, which has been reported to play a major role in both fear memory formation and fear expression [13, 24–29].

First, to develop a model to predict defense responses, a model was constructed based on EEG, ECG, and eye-tracking data that distinguishes the following two states during fear conditioning: (i) when a defense response occurs, and (ii) when a safety signal is provided. Because neural activity is high-dimensional and occurs in multiple locations, it is difficult to directly understand information related to defense responses. This issue can be addressed with machine learning, which was used to analyze hidden relationships between neural activity and human responses. To predict the frequency of defense responses, a learning model was adopted to classify medial prefrontal cortex (mPFC) neural activity, physiological response data for CRs, and controlled responses in a fear conditioning experiment.

Second, this model was shown to effectively measures the frequency of defense responses in a comprehensive situation that generates defense responses. Fearful and non-fearful short movie clips [30] were presented to generate defense responses. While watching each movie clip, physiological signals were measured. Defense responses were predicted in the time domain using the developed model. To validate the predicted result, participant’s emotional states that are known to correlate with defense responses in previous studies were measured. After watching each movie clip, the participant’s emotional state was determined by answering two questions that measure the participant’s subjective fear awareness and predictability/unpredictability of the watched movie clip’s contents. The latter represents the uncertainty of the existence of threat in the movie clip, which would cause a defense response [31].

Predicted defense response frequencies were compared with subjective fear ratings and the predictability/unpredictability of each clip. Subsequently, the results were compared with previous studies for which our model makes a valid prediction. It was confirmed that our model could be useful for understanding clinical studies of mood and anxiety disorders. Notably, the entire experiment was performed in a head-mounted virtual reality environment, which has recently been used in the neuroscience field to provide a high level of immersion for participants [32–36].

The primary aim of the study was to develop an effective model of temporal defense response elicitation that uses data from physiological signals (EEG, ECG, and eye tracking).

## Materials and methods

This study was approved by the institutional review board of Looxid Labs. Participants received information about the experiment and provided informed consent prior to participation.

### Experiment setup

Twelve adults were recruited (female, *n* = 3; male, *n* = 9; mean age (SD), 30.5 (3.77) years). All participants were right-handed, had no knowledge of neuropathology, and were familiar with virtual reality environments.

Before the experiment started, participants were provided explicit instructions about the purpose and overall procedures of the experiment, as suggested by guidelines for fear conditioning studies [8]. Participants sat on a chair in a relaxed position, with both hands on the armrests. The temperature of the experiment room was set at 20°C with an air conditioner to provide an immersed environment and to allow the collection of high-quality physiological data.

Physical features such as luminance and spatial frequency of emotion-related image stimuli are known to affect early brain activity [37, 38]. To minimize the effect of physical factors on brain activity, the mean frame pixel intensity of each task scene (mean (SD), 34.67 (3.83) pixels) and the sizes of the stimuli were set at similar levels.

### Procedures

The entire experiment consisted of three main parts. The first and second parts comprised stimulus habituation and conditioned fear acquisition to elicit defense responses represented by CRs. The final part comprised watching movie clips that were designed to elicit specific emotions (Fig 1). Tasks were conducted in a virtual reality environment within a mobile-based head-mounted headset (LooxidVR, Looxid Labs, Inc., CA, United States; Google Pixel phone, Google, Inc., CA, United States) with noise-cancellation headphones (Bose Q-20, Germany) and were performed sequentially without delay. The experimental setup for the fear conditioning task was constructed with reference to the method of Lau et al. [39].

**Fig 1.**
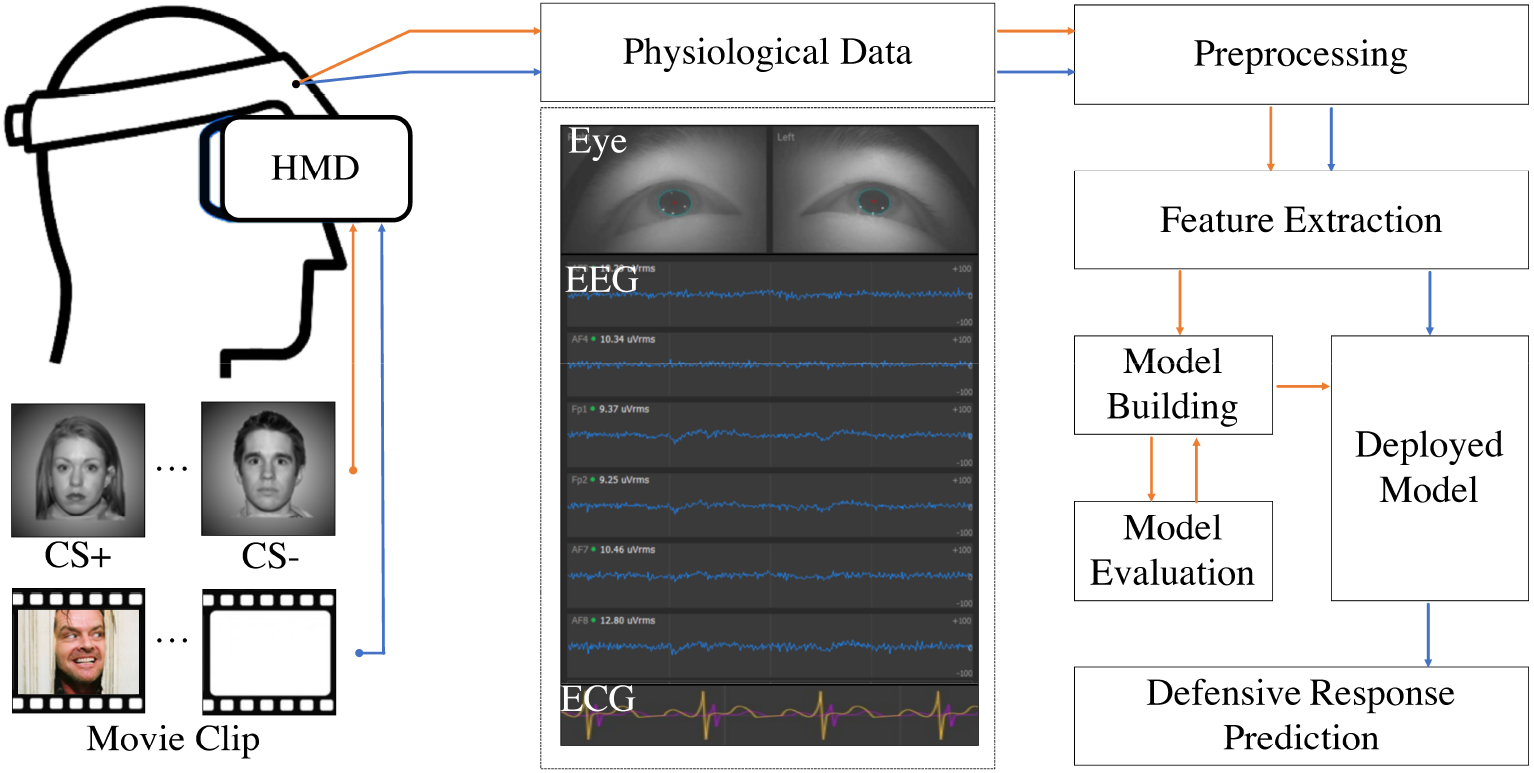
Schematic view of capturing and predicting defense responses with multi-modality data analysis.

In the habituation task (Fig 2B), five CS+, five controlled stimuli (CS-), and 10 auditory startle probes were presented in pseudorandom order without the US. The CS+ and CS- were both neutral human faces (NimStim [40]; 16 F, 32 M) and were each presented for 6 sec. To reduce the cognitive load, participants were able to easily discriminate between the CS+ (a neutral female face) and the CS- (a neutral male face). Inter-trial intervals lasted 10 to 12 sec between the offset of the prior CS and the onset of the next CS. To elicit a startle reflex at 50 ms, an auditory startle probe was presented in the form of a 105 dB burst of white noise (Fig 2A).

**Fig 2.**
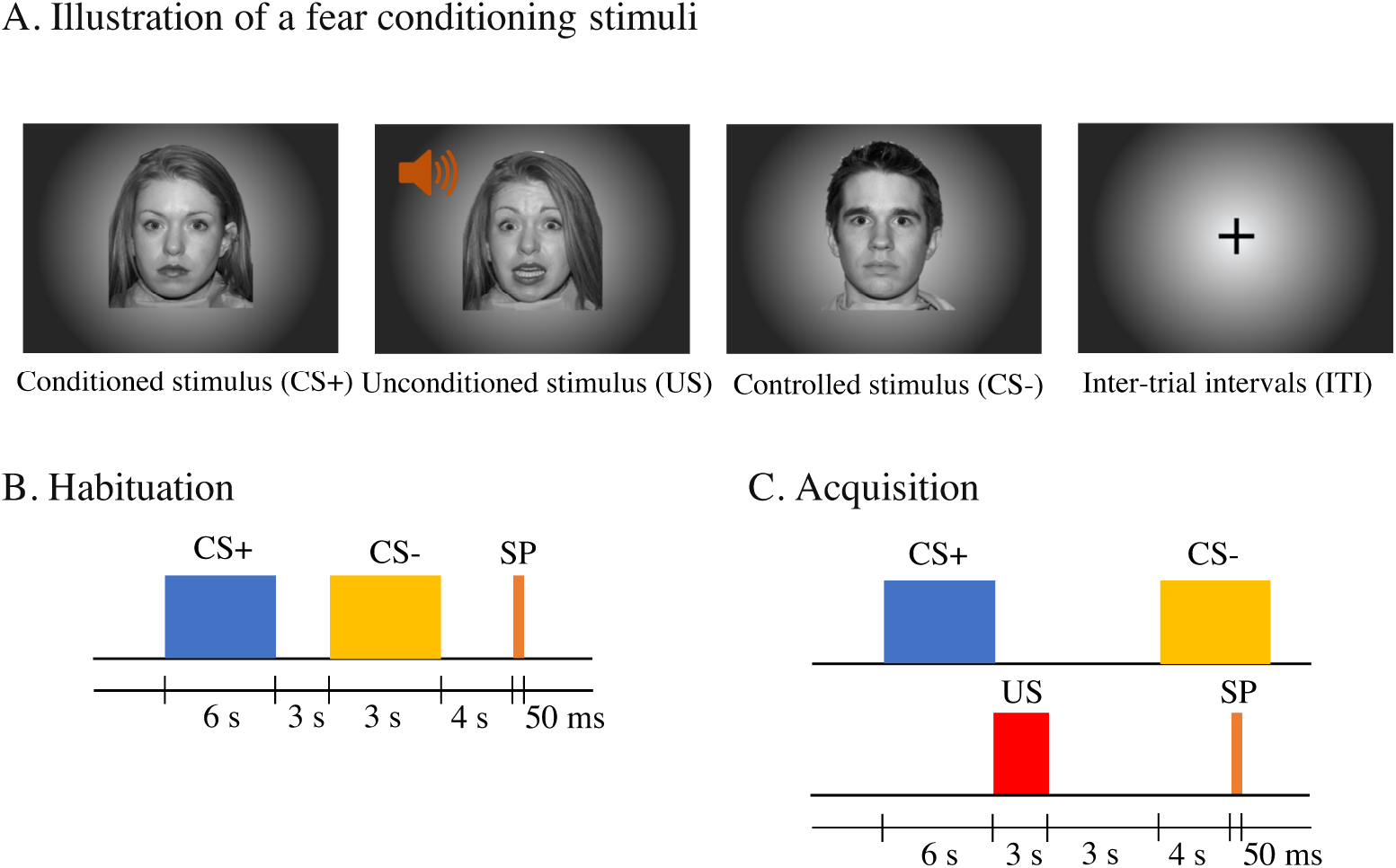
Fear conditioning procedures. (A) Illustration of each fear conditioning stimulus. A screaming female voice and fearful face image were provided as US. (B) Five CS+, five CS-, and 10 auditory startle probes (SP) were provided during the habituation process for each stimulus. (C) Ten acquisition trials with pseudorandomized CS+ and CS- were conducted.

In the fear acquisition task (Fig 2C), the US was presented immediately after CS+ offset with a 100% reinforcement rate but was not presented in the CS- condition. The US was a fearful female face with a loud female scream and was presented for 3 sec. The female face in the US was the same actress as in the CS+. The US scream was presented at approximately 80 dB for 1 sec at the onset of the US. During the acquisition training task, 10 CS+ with US and 10 CS- were presented in a pseudorandom order, with a single condition presented for no more than two consecutive trials. FPS and self-report questions were conducted to check whether fear conditioning was successful. Auditory startle probes were provided 4.5 sec after stimulus onset for the middle (5th) and last (10th) stimuli of each CS+/− acquisition trial.

Although FPS has been used to evaluate fear conditioning contingencies, this can be rendered difficult by the frequency of blinking or frowning, which can affect Electrooculography (EOG) signal quality. More frequent insertion of FPS might be an alternative solution. However, [22] suggested that frequent inclusion of FPS could affect the acquisition of conditioned fear. In addition, in our experiment, the FPS was not measured in some trials, usually because participants were blinking just before the stimuli were presented. These data were excluded from the FPS analysis according to published recommendations [41].

To overcome this limitation, after the acquisition training task, online self-report of fear/anxiety for the CS+/CS-/US and CS+/US contingency was performed. Participants were asked to rate on a ten-point Likert scale how fearful or anxious they felt in response to each CS+ or CS- neutral face image and to the US fearful female face image. Additionally, participants were asked, on a scale of 1 to 5, whether they knew when they were going to receive the US.

After the fear acquisition training task, eight movie clips inducing either fearful or tender emotions (FilmStim [30]) were presented to participants in a pseudorandom order (Table 1). Four of the eight clips elicited fear, and the other four elicited tenderness. Each movie clip was presented for 2–5 min. At the end of each clip, participants responded to two questions: 1) how much fear/anxiety did you feel in response to the movie clip; and 2) choose one of the following three statements: “When the movie clip was played, I recognized the genre and the detailed upcoming story because I have seen it before”, “When the movie clip was played, I recognized the genre and the upcoming story roughly because I have seen it before but could not remember the detailed story”, or “When the movie clip was played, I did not know the genre and the upcoming story at all because have not seen it before.” Participants were asked to rest sufficiently to allow recovery of their emotional state before starting the next movie clip.

**Table 1.** List of emotional movie clips provided in the experiment

| Index | Movie title | Duration | Pre-labeled genre |
| --- | --- | --- | --- |
| 1 | Ghost | 121 sec | Tenderness |
| 2 | The shining | 265 sec | Fearful |
| 3 | The exorcist | 105 sec | Fearful |
| 4 | Benny and Joone | 127 sec | Tenderness |
| 5 | Chucky II | 68 sec | Fearful |
| 6 | When a man loves a woman | 101 sec | Tenderness |
| 7 | Forrest Gump | 125 sec | Tenderness |
| 8 | The blair witch project | 239 sec | Fearful |

### Analysis

All analysis processes, including model development, training, and statistical analysis, were conducted with various functions in MATLAB 2017b. One-way analysis of variance (ANOVA) was used for all statistical analyses. For signal preprocessing and feature extraction, MATLAB codes provided by original research authors were used if they existed.

### Signal recordings and preprocessing

EEG signals from the medial prefrontal cortex region at Fp1, Fp2, AF3, AF4, AF7, and AF8 of the international 10-20 system were recorded with the LooxidVR signal acquisition system. Ground and reference nodes were located on the backs of both earlobes. EEG signals were amplified and digitized at a sampling rate of 250 Hz. These signals were notch filtered at 60 Hz and bandpass infinite impulse response (IIR) filtered in a range of 0.1–50 Hz. Jumping (jittering) noise was reduced with a technique reported by Patel et al. [42], and eyeblink noise was reduced with independent component analysis [43].

Vertical electrooculography (vEOG) signals from under the left or right eye were recorded at a sampling rate of 250 Hz to measure the startle response. As per the methods of Blumenthal et al. [41], participants were asked to identify their dominant hand and vEOG of the opposite eye was measured. This signal was notch filtered at 60 Hz and high-pass IIR filtered with a high-pass cutoff of 28 Hz. The signal was rectified and low-pass FIR was performed with a low-pass cutoff of 40 Hz. The peak signal amplitude 40 ms after startle probe onset was designated the startle response.

ECG signals of the wrist of the non-dominant hand were recorded at a sampling rate of 250 Hz. This signal was notch filtered at 60 Hz and bandpass IIR filtered in a range of 5–32 Hz. Two dimensional eye positions and pupil diameters of both eyes were measured with LooxidVR with two embedded eye-tracking cameras, at a sampling rate of 60 Hz.

EEG, EOG, ECG, and pupillary response signals were recorded and synchronized with LooxidVR with conductive adhesive hydrogel surface electrodes (KendallTM 100 Series Foam Electrodes, Covidien, Massachusetts, USA) and two embedded eye-tracking cameras.

### Model development

Features from EEG, pupillary response, and ECG signals for the last five CS+ and CS- acquisition trials in the conditioning acquisition step and the emotional-video-watching step were extracted for training and testing each classification feature model. First, EEG and physiological signals were collected for the first 300 ms (window size 2000 ms, step length 100 ms) after the onset of the last five CS+ or CS- acquisition trials, and used as input sample data for model training. From the raw data, 42 feature sets were extracted from 16 distinct types of features in the time, frequency, and time-frequency domains of the EEG and physiological signals. Detailed information about each feature set is presented in the Supplementary Information.

Second, a classification model was developed for each feature set. Before training each model, feature selection [44] was performed for each feature set to reduce the dimensionality. Subsequently, each feature model was trained from the reduced set of features. A Gaussian support vector machine (SVM) was used for the training model. The L2 regularization parameter (lambda) and kernel size (sigma) were estimated with the Bayesian optimization method [45]. Ten-fold cross-validation was applied, and the average fold loss was used to demonstrate model fitness.

## Result

### Development of defense response classification model

A defense response classification model was developed using sample data collected from the fear conditioning experiment. Sample data were validated with the validation method commonly used in fear conditioning experiments and then a classification model was trained with the collected sample data.

Before developing the classification model, it was validated whether the sample data properly represented labeled information. Checking the contingency between the CS+ and US conditions for each participant during acquisition was an important process from the perspective of data validation. Therefore, FPS and self-report measures were measured to validate the CS-US contingency and subjective fear ratings.

Specifically, EOG-based FPS in the first half of acquisition and the second half of acquisition after CS+ and CS- were measured. For all participants, the mean FPS amplitude differed significantly between the CS+ and CS- stimuli (Fig 3A). Additionally, the FPS for the first and second halves of the CS+ and CS- conditions were compared. The average amplitude of CS+ in the second half was lower than that during the first half, but still showed a meaningful difference from the CS- values in both halves (Supplementary Fig 1).

**Fig 3.**
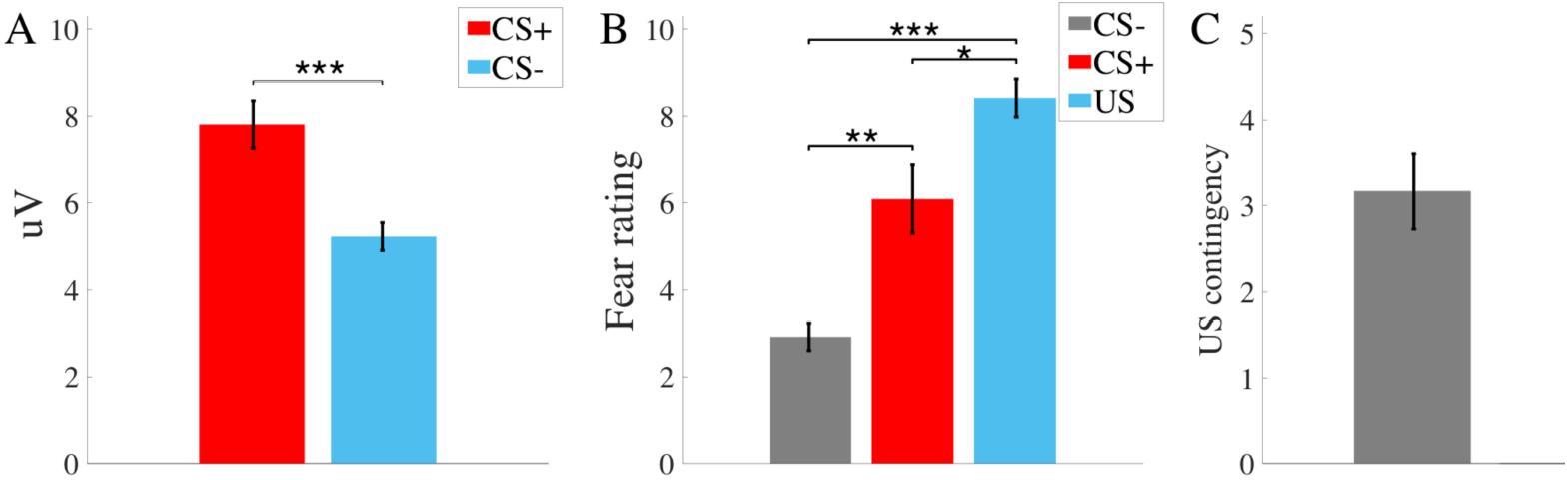
(A) Comparison of fear-potentiated startle response (FPS) for conditioned stimulus (CS+) and controlled stimulus (CS-). (Mean SEM, n = 15, 15). Trials for which the FPS could not be measured were excluded. (B) Comparison of subjective fear ratings for CS-, CS+ and unconditioned stimulus (US). (Mean ± SEM, n = 12). (C) Self-report responses for CS+–US contingency awareness. (Mean ± SEM, n = 12). ∗; p < 0.05, ∗∗; p < 0.005, ∗ ∗ ∗; p < 0.0005.

Moreover, participants completed four questionnaires after fear conditioning regarding subjective fear ratings (1 to 10) for each CS+/CS-/US stimulus and the contingency between the CS+ and US (1 to 5). Significant differences were observed between fear ratings for the different types of stimuli (Fig 3B). These results indicate that the conditioned fear acquisition was performed properly. Additionally, 4 participants who did not recognize the contingency between the CS+ and the US (correct contingencies lower than 3 out of 5) were excluded from further analysis (Fig 3C).

After validating the collected sample data from the fear conditioning experiment, defense response classification models were trained for each of the 42 extracted feature sets. The average cross-validation k-fold loss for all feature models between participants was 0.2210 (SEM, 0.0220). Among all feature models, the average band power feature model had the smallest mean k-fold loss (0.0460; SEM, 0.0160). The higher-order crossings (HOC) feature model for prediction of CRs had the second smallest mean k-fold loss and the smallest SEM (0.0462; SEM, 0.0077). Therefore, it was chosen for comparison with other high-ranked feature models (Fig 4). HOC measures the zero-crossing counts of the filtered signal, which represents the oscillation pattern of the signal in the time-series domain when a specific designed filter is applied. In the current study, a simple HOC system, which sequentially applied the backward difference operator as a high-pass filter was adopted [46, 47]. Physiological signals such as heart rate variability (HRV) and pupil diameter change are known as reliable features for fear conditioning. However, in our study, most physiological signal-based feature sets showed low model fitness. It is possible that this discrepancy is because of the short sample window (2000 ms), as autonomic physiological responses have slow onset and recovery compared with neural responses.

**Fig 4.**
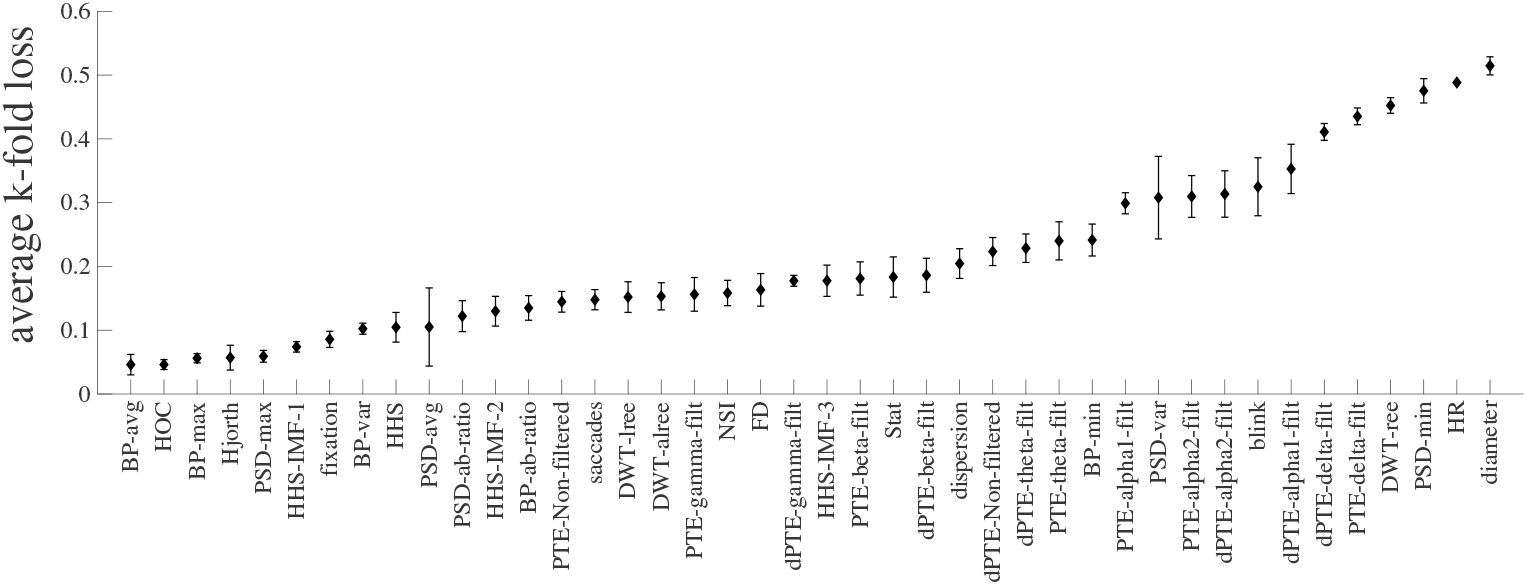
Average cross-validation k-fold loss for all feature sets between participants (Mean ± SEM, *n* = 8).

### Validation of defense response classification model

Next, the model was validated by examining the accuracy of predicted defense response occurrence ratios in a comprehensive environment that potentially generates defense responses. Eight short movie clips that elicited emotions of fear or tenderness were presented to participants. While participants watched each movie clip, defense response elicitation, implied by the frequency of occurrence of neural and physiological patterns similar to CRs, was predicted by the selected model every 100 ms, with 2000 ms windows. Predicted defense response incidence ratios for each movie clip were used as an index for how many neural responses or bodily responses similar to a defense response for visual and auditory stimuli were elicited while participants watched each emotional movie clip.

To validate predicted results, participants were asked to answer two self-report questions that are known to be related to defense response elicitation in previous studies at the end of each movie clip (as described above). For the first question, which measured subjective fear, the average rating for all movie scenes was 4.0417 (SEM, 0.8992). Moreover, the average ratings for horror and non-horror movie clips (as previously categorized by [30] were 6.7222 (SEM, 0.6196) and 1.3611 (SEM, 0.1566), respectively. Subjective fear ratings differed significantly between the two groups (p < 10^−17^). The second question consisted of three choices representing the unpredictability of threat being present in the movie clip, representing the unconscious threat defense system. For all movie clips, the proportions of participants choosing the most predictable, moderately predictable, least predictable options were 0.333, 0.208, and 0.458, respectively. In our analysis, choices 1 and 2, in which the subject had seen the movie before, were grouped (0.541) and compared with choice 3, in which the subject had not seen it before.

Based on the two self-report questions, participants’ experiences of the movie content could be classified into four emotional categories: 1) conscious fear with predictable threat context; 2) conscious fear with unpredictable threat context; 3) no conscious fear with predictably safe context; and 4) no conscious fear but unpredictably safe context.

Before comparing the predicted defense response ratios between the emotional categories, movie contents were compared based on three broader criteria. First, emotional content of the movie clips was classified as non-horror versus horror. Second, the movie clips were divided into fearful (subjective fear rating ≥ 5) versus non-fearful (rating < 5) based on the subjective fear ratings for each clip as rated by the participants. Third, the movie clips were divided into those that participants had seen before and those that they had not, regardless of the subjective fear ratings or pre-identified emotional genre. Each criterion represents whether participants felt conscious fear (criteria 1, 2) and the predictability of threat and safety in a context that causes defense responses and anxiety (criterion 3). Subsequently, we compared the predicted defense response occurrence ratio for the movie contents in the groups for each criterion.

For criteria 1 and 2, no statistically significant results were observed (First criteria, F_1,62_ = 0.2079, p = 0.6499; Second Criteria, F_1,62_ = 1.8269, p = 0.1814). However, for criterion 3, there was a statistically significant difference in the average predicted defense response occurrence ratio between participants that had seen the movies before and those who had not (F_1,62_ = 15.0593, p < 0.0005) (Figure 5A, 5B, 5C).

**Fig 5.**
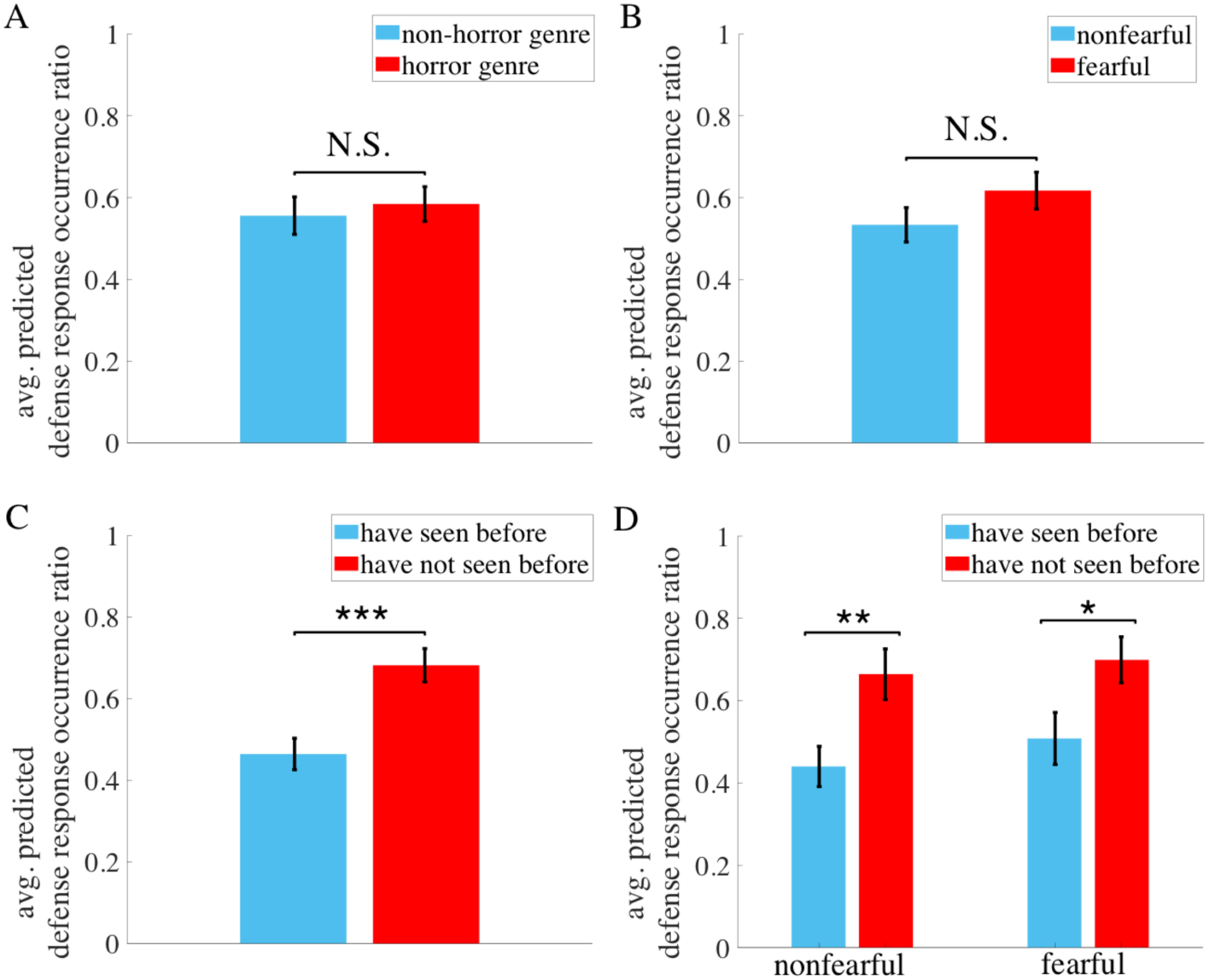
Comparison of average predicted defense response occurrence ratios between conditions (Mean ± SEM). (A) Group comparison based on prelabeled genres of movie clips (horror or non-horror) (*n* = 32,32). (B) Group comparison based on subjective fear ratings for each movie clip (n = 36,28). Non-fearful: fear rating < 5, Fearful: fear rating ≥ 5. (C) Group comparison based on previous experience of watching each movie clip (*n* = 33,31). (D) Group comparison between emotional categories measured by the two self-report questions. (nonfearful group, *n* = 21,15; fearful group, *n* = 12,16). N.S.; not significant. ∗; p < 0.05, ∗∗; p < 0.01, ∗ ∗ ∗; p < 0.0005.

Next, we compared the four emotional categories by combining the two self-report questions (Fig 5D). First, when participants felt no conscious fear, there was a significant difference depending on whether participants were able to predict the context of movie clips (F_1,34_ = 8.3375, p < 0.01). When participants felt conscious fear, a difference was also observed (rating ≥ 5) (F_1,26_ = 5.1080, p < 0.05). However, when participants were able to predict the context of the movie clip, no significant defense response ratio difference was observed between those who felt conscious fear and those who did not (F_1,31_ = 0.7169, p = 0.4036). The same trend was observed when participants were not able to predict the context of the movie clip (F_1,29_ = 0.1783, p = 0.6758).

## Discussion

Using a novel data-driven approach, we developed a model to predict temporal occurrence of defense responses. Previous methods of measuring defense responses include surveys, behavioral observation, FPS, and physiological observations. However, these methods measure a defense response at a particular timepoint, but it remains challenging to observe defense responses temporally. In particular, this limitation is a major challenge to understanding the relationship between anxiety and defense responses.

Anxiety is caused by chronic elicitation of defense responses to potential threat signals under conditions of uncertainty about the existence of an objective source of danger [7, 48–52]. Understanding the relationship between defense responses and anxiety is essential for studying clinical anxiety disorders such as GAD. As such, a method to continuously capture defense responses in the temporal domain is needed. Therefore, in this study, we adopted a model-based approach using fear conditioning to overcome such limitations.

We trained a binary classifier to classify mPFC neural activity, and physiological responses to CRs and controlled responses from fear conditioning to predict defense responses. Various features that have previously been reported to be related to emotions were extracted from mPFC EEG, ECG, and pupillary responses data. As a next step, we validated our model by comparing the predicted defense response occurrence ratio while participants watched eight emotional movie clips that provide fearful and nonfearful emotional states. For each movie clip, we measured two emotional situations, fear awareness and threat existence predictability. Because emotional situations are known to correlate with defense responses, we compared the predicted defense response ratio between the different situations.

In this study, there was no difference in predicted defense response ratios between participants who felt conscious fear and those who did not. In contrast, we observed that when the participant had not seen the movie clip before (representing unpredictability of threat existence), the predicted defense response ratio was much higher than for participants who had seen it before. This represents the predictability of threat existence, regardless of genre or awareness of fear. A significant difference in predicted defense response ratios between situations in which threat is predictable and those in which it is unpredictable was observed both when participants felt fear and when they did not. This suggests that mPFC neural activity patterns underlying defense responses against upcoming threats is related to threat existence predictability but not fear awareness.

Several previous studies have suggested that unpredictable threat maintains defense responses and thus increases anxiety [51, 53]. [53] reported that contextual fear induced by a US-unpaired CS was associated with a high startle amplitude measured by the FPS not only after CS onset, but also during the inter-trial interval, compared with that measured after a neutral and US-paired CS. Moreover, [51] suggested that unpredictability could lead to a sustained level of anxiety for sufficiently aversive stimuli. Our result is consistent with previous studies showing that defense responses occur more in situations with unpredictable threat than in those with predictable threat.

The existence of an unpredictable future threat arouses anxiety and makes individuals defensive against future situations [31]. In our experiment, without prior experience, participants were unable to know whether a movie clip was part of a love story or a horror movie. As a result, they remained defensive against any auditory, visual, or contextual indicators of potential threat that they had learned to be precursors of danger from experience or evolution [1]. Additionally, they paid attention to the consequences of each indicator to determine whether they were in a safe situation or not. Finally, when the movie ended, they were able to determine whether it had contained an actual threat or not, based on the collected information. Conscious fear would only occur with the actual presence of a threat or a memory of the consequences of specific indicators that they had learned before. For example, when participants watched non-horror movies without prior information, they unconsciously focused on possible indicators, and defense responses were elicited. However, without the actual appearance of a scary scene or a previously learned harmful indicator, no conscious fear would arise. Our study also shows discrepancy between conscious fear awareness and defense response occurrence.

LeDoux [4, 54] proposed a model to explain this discrepancy, claiming that fear can be described by a two-system model that comprises a conscious cognitive circuit and a unconscious defense survival circuit. Cognitive circuits mainly control conscious fearful feelings. Meanwhile, the defense survival circuit focuses on controlling unconscious threat defense responses against potential harmful situations and supporting the cognitive circuit. It is suggested that conscious fearful feelings occur without invoking the defense survival circuit and vice versa. Thus, comparison with previous studies suggests that our model predicts temporal occurrence of defense responses well.

## Conclusion

In summary, we report a novel data-driven approach using fear conditioning to predict temporal occurrence of defense responses. To validate our model, we compared defense response ratios in situations with differing fear awareness and threat existence predictability, which indicated that our model shows similar results to those in previous studies. Moreover, defense response ratios predicted by our model are consistent with previous studies using other methods such as FPS and self-report. Our study provides insight into measuring temporal defense responses in a comprehensive situation with threat and fear. This can help us to understand anxiety and related clinical disorders. To further improve the model, model stabilization can be conducted with a fear conditioning procedure that would enable the collection of more trial sample data.

## Supporting information

**S1 Fig.**
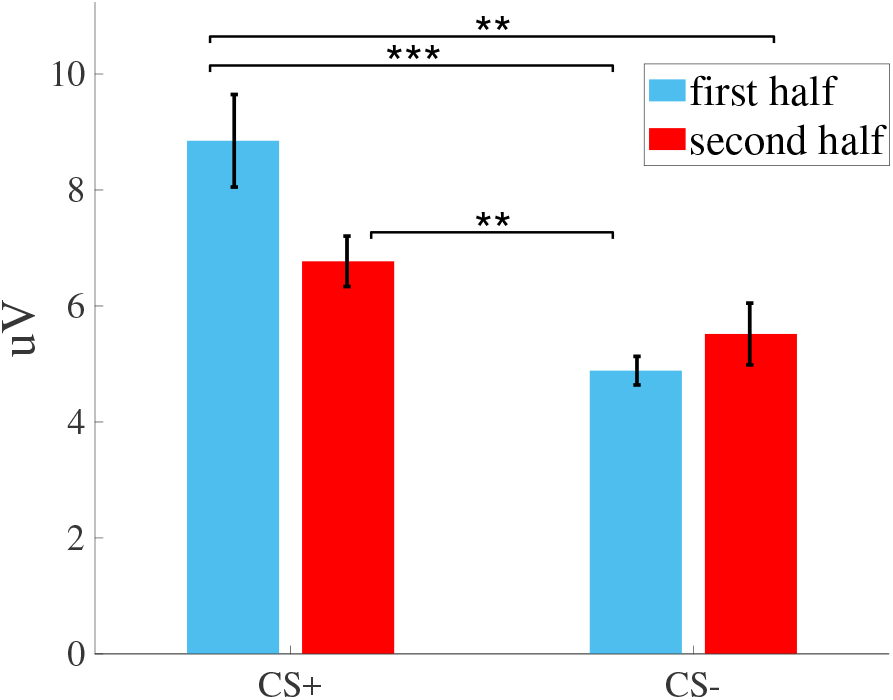
Differences in the fear-potentiated startle reflex (FPS) between the first and second halves of the CS+ and CS- sets (Mean ± SEM). One-way ANOVA was used for statistical analysis. **; p < 0.01, ***; p < 0.005

**S1 Appendix.**

### Feature extractions

Features were extracted from EEG, ECG signals and pupillary response. From 16 types of features, 42 distinct feature sets were constructed to train each binary classifier.

### EEG features

Various types of features from EEG signals that are relevant to the emotional index in time and frequency domain were extracted and analyzed. Details of the formula and information have been clearly described previously [46, 55–62].

#### Statistics

A statistical approach for analysis of physiological signals was first proposed by Picard et al. [56], and this concept has been adopted for many analyses of EEG recordings. In this experiment, power, mean, standard deviation, 1st difference, 2nd difference and normalization of the 1st difference and the 2nd difference of the EEG signal were measured and grouped as the ‘Stat’ feature set.

#### Hjorth

Hjorth developed 3 different time-domain EEG features that describe brain activity [57]. In this experiment, three parameters – activity, mobility and complexity – were measured and grouped as the ‘Hjorth’ feature set.

#### Non-stationary index (NSI)

The NSI represents the local average changes over time, independent of the magnitude fluctuation. Kroupi et al. [63], adapted the NSI to analyze emotional states using EEG. In this experiment, normalized NSIs were measured and grouped as the ‘NSI’ feature set.

#### Fractal dimension (FD)

The fractal dimension provides a statistical index of complexity that can be estimated through several methods. This experiment used Higuchi’s method [59], which is widely used to provide more accurate results than other methods, and these data were grouped as the ‘FD’ feature set.

#### Higher order crossings (HOC)

Petrantonakis and Hadjileontiadis [46], proposed the HOC-based emotion recognition system using EEG, which was originally introduced by Kedem. HOC measures the zero-crossing count for each filtered signal. In this experiment, simple HOC, which use the difference operator as a high-pass filter, for k = 1 to 20, were measured and grouped as the ‘HOC’ feature set.

#### Band power

One of the best-known frequency domain features for EEG recordings is band power features from different frequency bands. Power features capture the amplitude of neural oscillations in specific frequency band ranges. To transform the EEG recordings to the frequency domain, a discrete fast Fourier transform (FFT) was applied. In this experiment, average, standard deviation, variance, minimum and maximum band power of delta (1-4 Hz), theta (4-8 Hz), alpha (8-12 Hz), beta (12-30 Hz) and gamma (30-45 Hz) frequency bands and the ratio of average alpha and beta bands were transformed by both FFT and power spectral density (PSD). Features of each frequency band were grouped as ‘BP-avg’, ‘BP-min’, ‘BP-max’, ‘BP-var’, and ‘BP-ad-ratio’ feature sets for FFT features and ‘PSD-avg’, ‘PSD-min’, ‘PSD-max’, ‘PSD-var’, and ‘PSD-ab-ratio’ feature sets for PSD features.

#### Hilbert-Huang Spectrum (HHS)

Huang et al. [60], proposed the Hilbert-Huang transform for using empirical mode decomposition (EMD) to decompose a signal into several intrinsic mode functions (IMFs). The instantaneous amplitude and instantaneous frequency of the IMFs were examined with Hilbert spectral analysis (HSA). In this experiment, the average amplitudes of the delta, theta, alpha, beta and gamma frequency bands were calculated and grouped as the ‘HHS’ feature set. In addition, the root mean square and maximum instantaneous amplitude, mean and weighted mean instantaneous frequency of each IMF, levels 1 to 3, were measured and grouped as the ‘HHS-IMF-1’, ‘HHS-IMF-2’, and ‘HHS-IMF-3’ feature sets according to [64].

#### Discrete wavelet transforms (DWT)

DWT refers to various wavelet transforms that decompose the signals into discretely sampled wavelets that capture different frequency and temporal information. Compared with the Fourier transform, DWT offers advantages for analyzing time-series signals, in that DWT basis functions are finite in time but the basis of the Fourier transform is not. Wavelet transforms decompose the signals into different approximations and levels of detail that represent different frequency ranges. Murugappan et al. [61] used a db4 wavelet transform to extract the energy-based features from fourth-level detail coefficients corresponding to alpha-band frequency of EEG recordings to classify emotions. In this experiment, the recursive energy efficiency (REE), logarithmic REE (LREE) and absolute logarithmic REE (ALREE) of each level of energy were measured. These features were grouped as the ‘DWT-ree’, ‘DWT-lree’, and ‘DWT-alree’ feature sets.

#### Phase transfer entropy (PTE)

PTE measures the energy-based information flow from the transfer of phase between different neuronal channels. This feature was recently introduced by Lobier et al. [65] and represent the directed connectivity networks estimated from neuronal oscillation patterns among EEG/MEG channels [62]. In our experiment, PTE and dPTE (normalized PTE) for the delta, theta, alpha1 (8 – 10 Hz), alpha2 (10 – 13 Hz), beta and gamma frequency ranges were measured as described by Hillebrand et al [62]. These features were grouped as the ‘PTE-non-filtered’, ‘dPTE-non-filtered’, ‘PTE-delta-filt’, ‘dPTE-delta-filt’, ‘PTE-theta-filt’, ‘dPTE-theta-filt’, ‘PTE-alpha1-filt’, ‘dPTE-alpha1-filt’, ‘PTE-alpha2-filt’, ‘dPTE-alpha2-filt’, ‘PTE-beta-filt’, ‘dPTE-beta-filt’, ‘PTE-gamma-filt’, and ‘dPTE-gamma-filt’ feature sets.

### Pupil features

From relative diameter and x, y position in eye-tracking camera images of both pupils, emotion-related features were extracted. Lu et al. [66] provides a detailed formula and description.

#### Blink

The average eye-blinking frequency and the mean and standard deviation of blinking duration were measured and grouped as the ‘blink’ feature set.

#### Pupil diameter

The mean and standard deviation of the pupil diameter and the PSD in four frequency ranges (0–0.2 Hz, 0.2–0.4 Hz, 0.4–0.6 Hz, 0.6–1 Hz) were measured and grouped as the ‘diameter’ feature set.

#### Dispersion, fixation

Mean and standard deviation of dispersion and fixation were measured and grouped as the ‘dispersion’ and ‘fixation’ feature sets.

#### Saccades

The mean and standard deviation of saccade duration and amplitude were measured and grouped as the ‘saccades’ feature set.

### ECG feature

ECG signals were recorded from the wrist opposite from the subject’s dominant hand.

#### Heart rate variability (HRV)

HRV was calculated as the average time interval of R peaks of the QRS complex and represented as the ‘HR’ feature set.

## Acknowledgments

We thank Kibum Choi and Hyeonyoung Choi for their help with developing the SW and HW environment for experiment. Also we thank Jaeho Bae for designing image of the fear conditioning experiment. This research was supported by the Looxid Labs. All data are stored on Looxid Labs’s server and are available upon request.

## Notes

### Competing Interest Statement

The authors have declared no competing interest.

## References

1. Blanchard DC, Blanchard RJ. .4 Defensive behaviors, fear, and anxiety. Handbook of behavioral neuroscience. 2008;17:63–79.

2. Blanchard RJ, Yudko EB, Rodgers RJ, Blanchard DC. Defense system psychopharmacology: an ethological approach to the pharmacology of fear and anxiety. Behavioural brain research. 1993;58(1-2):155–165.

3. Miles L, Davis M, Walker D. Phasic and sustained fear are pharmacologically dissociable in rats. Neuropsychopharmacology. 2011;36(8):1563.

4. LeDoux JE. Coming to terms with fear. Proc Natl Acad Sci. 2014;111(8):2871–2878.

5. Mineka S, Zinbarg R. A contemporary learning theory perspective on the etiology of anxiety disorders: it’s not what you thought it was. American psychologist. 2006;61(1):10.

6. Foa EB, Zinbarg R, Rothbaum BO. Uncontrollability and unpredictability in post-traumatic stress disorder: an animal model. Psychological bulletin. 1992;112(2):218.

7. Lang PJ, Davis M, Öhman A. Fear and anxiety: animal models and human cognitive psychophysiology. Journal of affective disorders. 2000;61(3):137–159.

8. Lonsdorf TB, Menz MM, Andreatta M, Fullana MA, Golkar A, Haaker J, et al. Don’t fear ‘fear conditioning’: Methodological considerations for the design and analysis of studies on human fear acquisition, extinction, and return of fear. Neurosci Biobehav Rev. 2017;77:247–285.

9. Blanchard DC. Stimulus, environmental, and pharmacological control of defensive behaviors. 1997;.

10. Karalis N, Dejean C, Chaudun F, Khoder S, Rozeske RR, Wurtz H, et al. 4-Hz oscillations synchronize prefrontal–amygdala circuits during fear behavior. Nat Neurosci. 2016;19(4):605.

11. Korn CW, Staib M, Tzovara A, Castegnetti G, Bach DR. A pupil size response model to assess fear learning. Psychophysiology. 2017;54(3):330–343.

12. Leuchs L, Schneider M, Czisch M, Spoormaker VI. Neural correlates of pupil dilation during human fear learning. Neuroimage. 2017;147:186–197.

13. Visser RM, Scholte HS, Beemsterboer T, Kindt M. Neural pattern similarity predicts long-term fear memory. Nature neuroscience. 2013;16(4):388.

14. Critchley HD. Electrodermal responses: what happens in the brain. The Neuroscientist. 2002;8(2):132–142.

15. Hamm AO, Vaitl D. Affective learning: Awareness and aversion. Psychophysiology. 1996;33(6):698–710.

16. Bradley MM, Codispoti M, Cuthbert BN, Lang PJ. Emotion and motivation I: defensive and appetitive reactions in picture processing. Emotion. 2001;1(3):276.

17. Jansen DM, Frijda NH. Modulation of the acoustic startle response by film-induced fear and sexual arousal. Psychophysiology. 1994;31(6):565–571.

18. Vrana SR, Spence EL, Lang PJ. The startle probe response: a new measure of emotion? Journal of abnormal psychology. 1988;97(4):487.

19. Davis M. Neural systems involved in fear and anxiety measured with fear-potentiated startle. American Psychologist. 2006;61(8):741.

20. Davis M, Falls WA, Campeau S, Kim M. Fear-potentiated startle: a neural and pharmacological analysis. Behavioural brain research. 1993;58(1-2):175–198.

21. LeDoux JE. Emotion circuits in the brain. Annu Rev Neurosci. 2000;23(1):155–184.

22. Sjouwerman R, Niehaus J, Kuhn M, Lonsdorf TB. Don’t startle me—interference of startle probe presentations and intermittent ratings with fear acquisition. Psychophysiology. 2016;53(12):1889–1899.

23. Chien J, Colloca L, Korzeniewska A, Cheng J, Campbell C, Hillis A, et al. Oscillatory EEG activity induced by conditioning stimuli during fear conditioning reflects Salience and Valence of these stimuli more than Expectancy. Neuroscience. 2017;346:81–93.

24. Gilmartin MR, Balderston NL, Helmstetter FJ. Prefrontal cortical regulation of fear learning. Trends Neurosci. 2014;37(8):455–464.

25. Dejean C, Courtin J, Karalis N, Chaudun F, Wurtz H, Bienvenu TC, et al. Prefrontal neuronal assemblies temporally control fear behaviour. Nature. 2016;535(7612):420.

26. Courtin J, Bienvenu T, Einarsson E, Herry C. Medial prefrontal cortex neuronal circuits in fear behavior. Neuroscience. 2013;240:219–242.

27. Sierra-Mercado D, Padilla-Coreano N, Quirk GJ. Dissociable roles of prelimbic and infralimbic cortices, ventral hippocampus, and basolateral amygdala in the expression and extinction of conditioned fear. Neuropsychopharmacology. 2011;36(2):529.

28. Vidal-Gonzalez I, Vidal-Gonzalez B, Rauch SL, Quirk GJ. Microstimulation reveals opposing influences of prelimbic and infralimbic cortex on the expression of conditioned fear. Learn Mem. 2006;13(6):728–733.

29. Kim MJ, Gee DG, Loucks RA, Davis FC, Whalen PJ. Anxiety dissociates dorsal and ventral medial prefrontal cortex functional connectivity with the amygdala at rest. Cereb Cortex. 2010;21(7):1667–1673.

30. Schaefer A, Nils F, Sanchez X, Philippot P. Assessing the effectiveness of a large database of emotion-eliciting films: A new tool for emotion researchers. Cogn Emotion. 2010;24(7):1153–1172.

31. Grupe DW, Nitschke JB. Uncertainty and anticipation in anxiety: an integrated neurobiological and psychological perspective. Nat Rev Neurosci. 2013;14(7):488.

32. Lin JHT. Fear in virtual reality (VR): Fear elements, coping reactions, immediate and next-day fright responses toward a survival horror zombie virtual reality game. Comput Hum Behav. 2017;72:350–361.

33. Reichenberger J, Porsch S, Wittmann J, Zimmermann V, Shiban Y. Social Fear Conditioning Paradigm in Virtual Reality: Social vs. Electrical Aversive Conditioning. Front Psychol. 2017;8:1979.

34. Glotzbach-Schoon E, Andreatta M, Reif A, Ewald H, Tröger C, Baumann C, et al. Contextual fear conditioning in virtual reality is affected by 5HTTLPR and NPSR1 polymorphisms: effects on fear-potentiated startle. Front Behav Neurosci. 2013;7:31.

35. Bohil CJ, Alicea B, Biocca FA. Virtual reality in neuroscience research and therapy. Nat Rev Neurosci. 2011;12(12):752.

36. Alvarez RP, Johnson L, Grillon C. Contextual-specificity of short-delay extinction in humans: renewal of fear-potentiated startle in a virtual environment. Learn Mem. 2007;14(4):247–253.

37. Alorda C, Serrano-Pedraza I, Campos-Bueno JJ, Sierra-Vázquez V, Montoya P. Low spatial frequency filtering modulates early brain processing of affective complex pictures. Neuropsychologia. 2007;45(14):3223–3233.

38. Mavratzakis A, Herbert C, Walla P. Emotional facial expressions evoke faster orienting responses, but weaker emotional responses at neural and behavioural levels compared to scenes: A simultaneous EEG and facial EMG study. Neuroimage. 2016;124:931–946.

39. Lau JY, Britton JC, Nelson EE, Angold A, Ernst M, Goldwin M, et al. Distinct neural signatures of threat learning in adolescents and adults. Proc Natl Acad Sci. 2011;108(11):4500–4505.

40. Tottenham N, Tanaka JW, Leon AC, McCarry T, Nurse M, Hare TA, et al. The NimStim set of facial expressions: judgments from untrained research participants. Psychiatry Res. 2009;168(3):242–249.

41. Blumenthal TD, Cuthbert BN, Filion DL, Hackley S, Lipp OV, Van Boxtel A. Committee report: Guidelines for human startle eyeblink electromyographic studies. Psychophysiology. 2005;42(1):1–15.

42. Patel AX, Bullmore ET. A wavelet-based estimator of the degrees of freedom in denoised fMRI time series for probabilistic testing of functional connectivity and brain graphs. NeuroImage. 2016;142:14–26.

43. Makeig S, Bell AJ, Jung TP, Sejnowski TJ. Independent component analysis of electroencephalographic data. In: Advances in neural information processing systems; 1996. p. 145–151.

44. Roffo G, Melzi S, Castellani U, Vinciarelli A. Infinite Latent Feature Selection: A Probabilistic Latent Graph-Based Ranking Approach. In: IEEE International Conference on Computer Vision; 2017.

45. Snoek J, Larochelle H, Adams RP. Practical bayesian optimization of machine learning algorithms. In: Advances in neural information processing systems; 2012. p. 2951–2959.

46. Petrantonakis PC, Hadjileontiadis LJ. Emotion recognition from EEG using higher order crossings. IEEE Transactions on Information Technology in Biomedicine. 2010;14(2):186–197.

47. Kedem B, Yakowitz S. Time series analysis by higher order crossings. IEEE press New York; 1994.

48. Barlow DH. Unraveling the mysteries of anxiety and its disorders from the perspective of emotion theory. American psychologist. 2000;55(11):1247.

49. Bouton ME, Mineka S, Barlow DH. A modern learning theory perspective on the etiology of panic disorder. Psychological review. 2001;108(1):4.

50. Grillon C. Startle reactivity and anxiety disorders: aversive conditioning, context, and neurobiology. Biological psychiatry. 2002;52(10):958–975.

51. Grillon C, Baas JP, Lissek S, Smith K, Milstein J. Anxious responses to predictable and unpredictable aversive events. Behavioral neuroscience. 2004;118(5):916.

52. Lake JI, LaBar KS. Unpredictability and uncertainty in anxiety: a new direction for emotional timing research. Frontiers in integrative neuroscience. 2011;5:55.

53. Vansteenwegen D, Iberico C, Vervliet B, Marescau V, Hermans D. Contextual fear induced by unpredictability in a human fear conditioning preparation is related to the chronic expectation of a threatening US. Biological psychology. 2008;77(1):39–46.

54. LeDoux JE, Pine DS. Using neuroscience to help understand fear and anxiety: a two-system framework. Am J Psychiatry. 2016;173(11):1083–1093.

55. Jenke R, Peer A, Buss M. Feature extraction and selection for emotion recognition from EEG. IEEE Transactions on Affective Computing. 2014;5(3):327–339.

56. Picard RW, Vyzas E, Healey J. Toward machine emotional intelligence: Analysis of affective physiological state. IEEE transactions on Pattern Analysis and Machine Intelligence. 2001;23(10):1175–1191.

57. Hjorth B. EEG analysis based on time domain properties. Electroencephalogr Clin Neurophysiol. 1970;29(3):306–310.

58. Kroupi E, Yazdani A, Ebrahimi T. EEG correlates of different emotional states elicited during watching music videos. In: Affect Comput Intell Interact. Springer; 2011. p. 457–466.

59. Higuchi T. Approach to an irregular time series on the basis of the fractal theory. Physica D: Nonlinear Phenomena. 1988;31(2):277–283.

60. Huang NE, Shen Z, Long SR, Wu MC, Shih HH, Zheng Q, et al. The empirical mode decomposition and the Hilbert spectrum for nonlinear and non-stationary time series analysis. In: Proc Roy Soc London Series A: Math, Phys Eng Sci. vol. 454. The Royal Society; 1998. p. 903–995.

61. Murugappan M, Ramachandran N, Sazali Y. Classification of human emotion from EEG using discrete wavelet transform. J Biomed Sci Eng. 2010;3(04):390.

62. Hillebrand A, Tewarie P, Van Dellen E, Yu M, Carbo EW, Douw L, et al. Direction of information flow in large-scale resting-state networks is frequency-dependent. Proc Natl Acad Sci. 2016;113(14):3867–3872.

63. Hausdorff JM, Lertratanakul A, Cudkowicz ME, Peterson AL, Kaliton D, Goldberger AL. Dynamic markers of altered gait rhythm in amyotrophic lateral sclerosis. J Appl Physiol. 2000;88(6):2045–2053.

64. Zong C, Chetouani M. Hilbert-Huang transform based physiological signals analysis for emotion recognition. In: Signal processing and information technology (isspit), 2009 ieee international symposium on. IEEE; 2009. p. 334–339.

65. Lobier M, Siebenhühner F, Palva S, Palva JM. Phase transfer entropy: a novel phase-based measure for directed connectivity in networks coupled by oscillatory interactions. Neuroimage. 2014;85:853–872.

66. Lu Y, Zheng WL, Li B, Lu BL. Combining Eye Movements and EEG to Enhance Emotion Recognition. In: IJCAI. vol. 15; 2015. p. 1170–1176.

